# An efficient eukaryotic cell-free expression and correlative cryo-electron tomography platform for structural biology of macromolecular complexes

**DOI:** 10.1101/2025.10.18.683253

**Authors:** Vikas Tillu, Dominic Hunter, Kai-En Chen, Jessica Smith, Olivia Nassar, James Rae, Emma Sierecki, Bostjan Kobe, Brett M. Collins, Yann Gambin, Robert G. Parton, Nicholas Ariotti

## Abstract

Cell-free expression using *Leishmania tarentolae* lysates allows rapid expression of eukaryotic proteins directly from DNA templates. We develop a pipeline that combines cell-free expression system with cryogenic fluorescence microscopy that we term CC-FLEXCET (Correlative Cell-Free *Leishmania* EXpression and Cryo-Electron Tomography), to target and visualize expressed protein complexes by cryo-electron tomography at high resolution. We demonstrate the utility of this method by structurally characterising the filaments of the full-length apoptosis-associated speck like protein containing CARD (ASC) protein. Cell-free expression of ASC results in a polymeric structure characteristic of its cellular speck assembly. Sub-tomogram averaging allows us to resolve both the pyrin domain (PYD) to medium resolution, and show, for the first time, the arrangement of the flexibly linked caspase recruitment domain (CARD). Finally, we observed an interaction between the ASC filament and the *L. tarentolae* ribosome. Using template matching and quantitative approaches, we characterise this interaction and determine that there is a random structural association between the filament and the ribosome, with 57% of ribosomes oriented with the LSU toward the ASC polymer. CC-FLEXCET facilitates structural analysis of macromolecules and protein-lipid assemblies without need of purification, providing a pipeline from DNA template to protein expression to cryo-tilt series acquisition, within a single day.

## Introduction

Recombinant expression systems are typically required for biochemical characterisation of protein structure and function. Advances in cryo-electron microscopy have driven a concurrent need for expression and purification of large and often heterogeneous macromolecular complexes for structure determination. Of the various recombinant approaches, *Escherichia coli* is the most ubiquitously utilised expression system which, while highly scalable, can suffer from poor protein folding capacity, especially with respect to eukaryotic proteins ^1^. While eukaryotic expression systems, such as mammalian human embryonic kidney 293 (HEK) and insect models such as Sf9 cells are generally superior with respect to eukaryotic protein folding, they are less scalable than *E. coli* and can be cost prohibitive^2^. For decades, cell-free expression systems have been used to study protein behaviour and cellular processes in a semi-pure state^3,4^, but their utility in structural biology is only just beginning to be realised.

Cell-free systems include rabbit reticulocyte lysate (RRL), wheat germ extract (WGE), insect cell extract (ICE) and, more recently, *Leishmania tarentolae* extract (LTE)^5,6^ . A major advantage of cell-free expression is that it can generate high-quality eukaryotic protein, while simultaneously facilitating functional expression of other proteins related to protein regulation. As such, these systems have been applied to study enzymatic functions, protein directed evolution, and translational mechanisms with great success ^3,4,7^. They have facilitated unique insights into post-translational modifications such as lipidation and phosphorylation ^5,8^, and we have previously used these methods to investigate and validate protein-protein interactions *in vitro* ^9^, describe and characterise the stoichiometries involved in protein-protein interactions ^10^ and validate phosphorylation of protein targets by co-expressing kinases ^8^. While cell-free systems are somewhat scalable ^3,4,7^, the volumes of lysate required for typical structural biology approaches and the costs involved in generating these reactions have proven limiting. Only a few successful examples of structures of proteins produced by cell-free systems have been published, including single particle cryoEM structures of GPCRs produced in *E. coli* lysates and reconstituted into nanodiscs^11,12^, and crystal and cryoEM structures of proteins produced in WGE^13–15^. A system that routinely facilitates rapid high quality eukaryotic protein expression, yet still allows for high-resolution structural analyses of macromolecular complexes without purification, would represent a major advance.

Recent developments in *in situ* cryo-electron tomography (cryo-ET) and sub-tomogram averaging are now driving structural biodiscovery in cells, and large and abundant protein structures such as the ribosome are being solved in the cellular cytoplasm to near atomic resolution^16^. These approaches can be highly successful when dealing with large molecular weight, well-characterised protein complexes like the ribosome; however, they are less successful when attempting structural analyses on smaller or poorly characterised proteins^17^. To perform these kinds of analyses, other methods are required to validate protein targets present in non-homogenous mixtures like the cytoplasm. These techniques can include template matching^18,19^, genetically expressed molecular markers^20^ and exogenous addition of signposts^21^. The most common approach, however, is the application of cryo-fluorescence imaging to correlate between fluorescently-labelled targets in a cryo-fluorescence microscope and the cryo-electron microscope (cryo-CLEM)^22,23^. These developments are now underpinning a change in the way we can perform structural biology.

Here, we sought to combine the advantages of cell-free protein production with cryo-CLEM and cryo-ET for protein structural studies. We describe the development and application CC-FLEXCET; Correlative Cell-Free *Leishmania* EXpression and Cryo-Electron Tomography. Using cryogenic correlative light and electron tomography, we can target large, fluorescently labelled macromolecular complexes expressed directly in the *Leishmania tarentolae* lysate. Such samples are well suited to high-resolution imaging and sub-tomogram averaging, to determine structures of our proteins of interest without biochemical purification. We demonstrate the system by determining the structures of endogenous *L. tarantolae* microtubules and ribosomes, by using CC-FLEXCET to characterise exogenously expressed proteins, and to provide the first structural analyses of the filamentous assembly of full-length human ASC at nanometre resolution.

## Results

### Structural analysis of *Leishmania tarentolae* promastigotes and cell-free lysates

To generate a system that will facilitate a rapid pipeline to express and structurally characterise proteins of interest directly in cell-free lysates, we first sought to establish a clear structural understanding of the starting material that is present in the *Leishmania tarentolae* extracts. Given the importance of apicomplexans and other *Leishmania* species as parasitic pathogens in human health numerous ultrastructural analyses have been conducted^24,25^, but there is a limited understanding of the ultrastructure of *L. tarentolae*. As such, we initially characterised the cultured promastigote form of these organisms using conventional transmission electron microscopy. We observed morphologies consistent with other *Leishmania* spp. and could resolve various sub-cellular structures including the prominent flagellum, the kinetoplast, and the subpellicular microtubular array (Fig. 1A-H).

**Figure 1.**
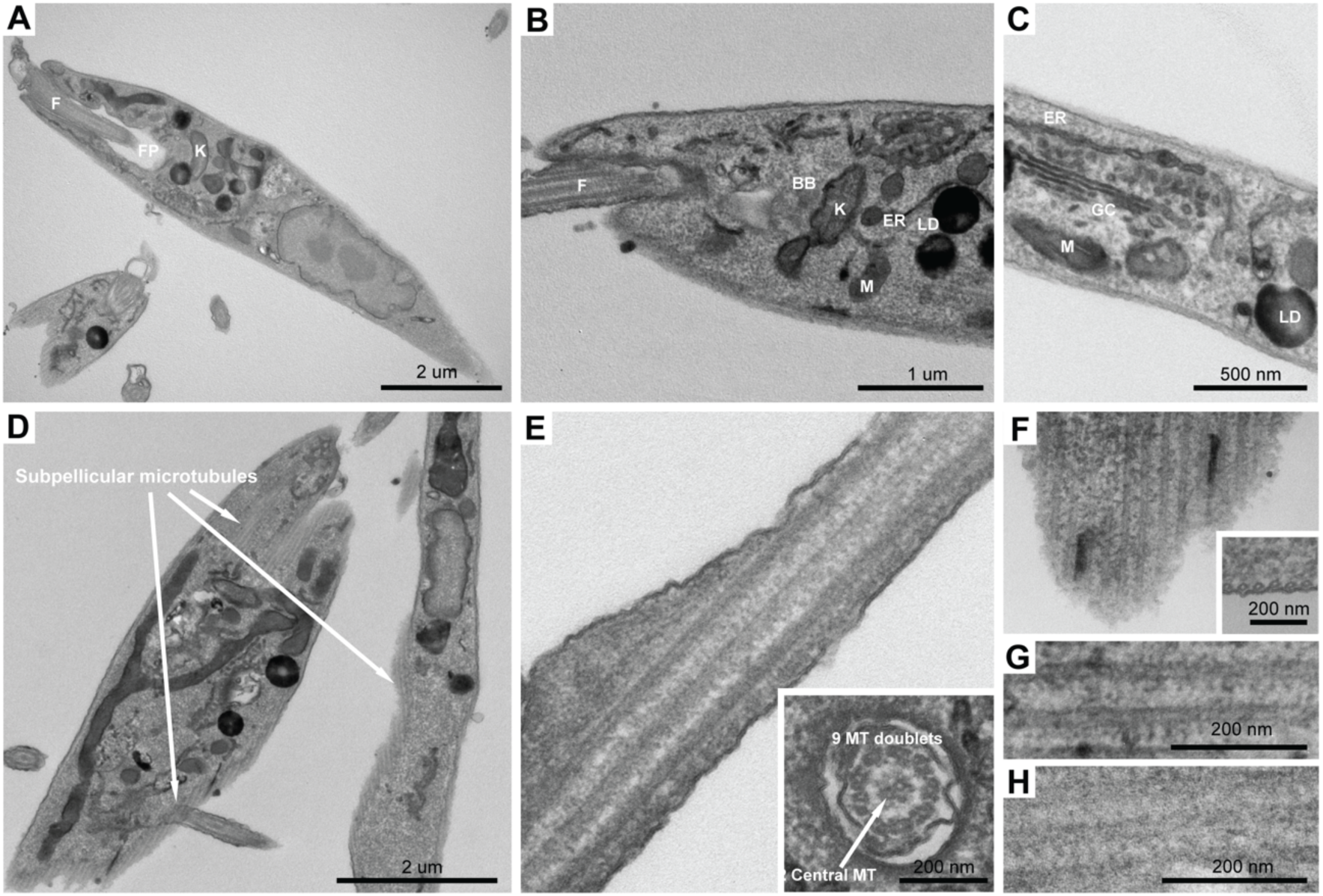
Ultrastructure of *Leishmania tarentolae* by transmission electron microscopy. (A) Whole-cell image of the promastigote form to *L. tarentolae,* showing the kinetoplast (K) arranged at the base of the flagellum (F) and the flagellar pocket (FP). (B) Higher-magnification image, showing the basal body (BB), flagellum (F), kinetoplast (K), lipid droplet (LD) and mitochondria (M). (C) High-magnification image, highlighting the organisation of endogenous membrane-containing compartments, including the endoplasmic reticulum (ER), mitochondria (M), Golgi complex (GC) and lipid droplets (LD). (D-E) The organisation and abundance of microtubules in *L. tarentolae* cells. The subpellicular microtubules (D,F-H) are abundant around the entire periphery of the cell and the organisation of the microtubules in the flagellum can be clearly distinguished in E and the inset.

We next examined the content of the cell-free LTE. Lysates were prepared as described previously (Fig. 2A) ^5^ and spotted to a transmission EM grid for negative staining. Within the bulk LTE, we observed significant biomolecular material including vesicles, microtubules, and actin fibres (Fig. 2B-D). With these samples, we performed an unbiased cryo-ET screen of the LTE, where we imaged random areas of the EM grids, including areas from thin (<100 nm) to thick vitrified ice (>500 nm) (Fig. 2E-I). As expected, the tomograms from thicker regions resulted in a lower success rate for reconstruction, at a 20% failure rate (Fig. 2J). The successfully reconstructed cryo-electron tomograms revealed many morphologically identifiable structures that included membrane vesicles (Fig. 2E-E’’), ribosomes (Fig. 2F, F’), actin filaments (Fig. 2G-G’’), microtubules (Fig. 2H), and whole lipid droplets (Fig. 2I). From these reconstructions, we observed that 90% of the cryo-electron tomograms contained membrane-bound vesicles, 75% ribosomes, 20% microtubules, 62% actin fibres and 6% lipid droplets (Fig. 2J).

**Figure 2.**
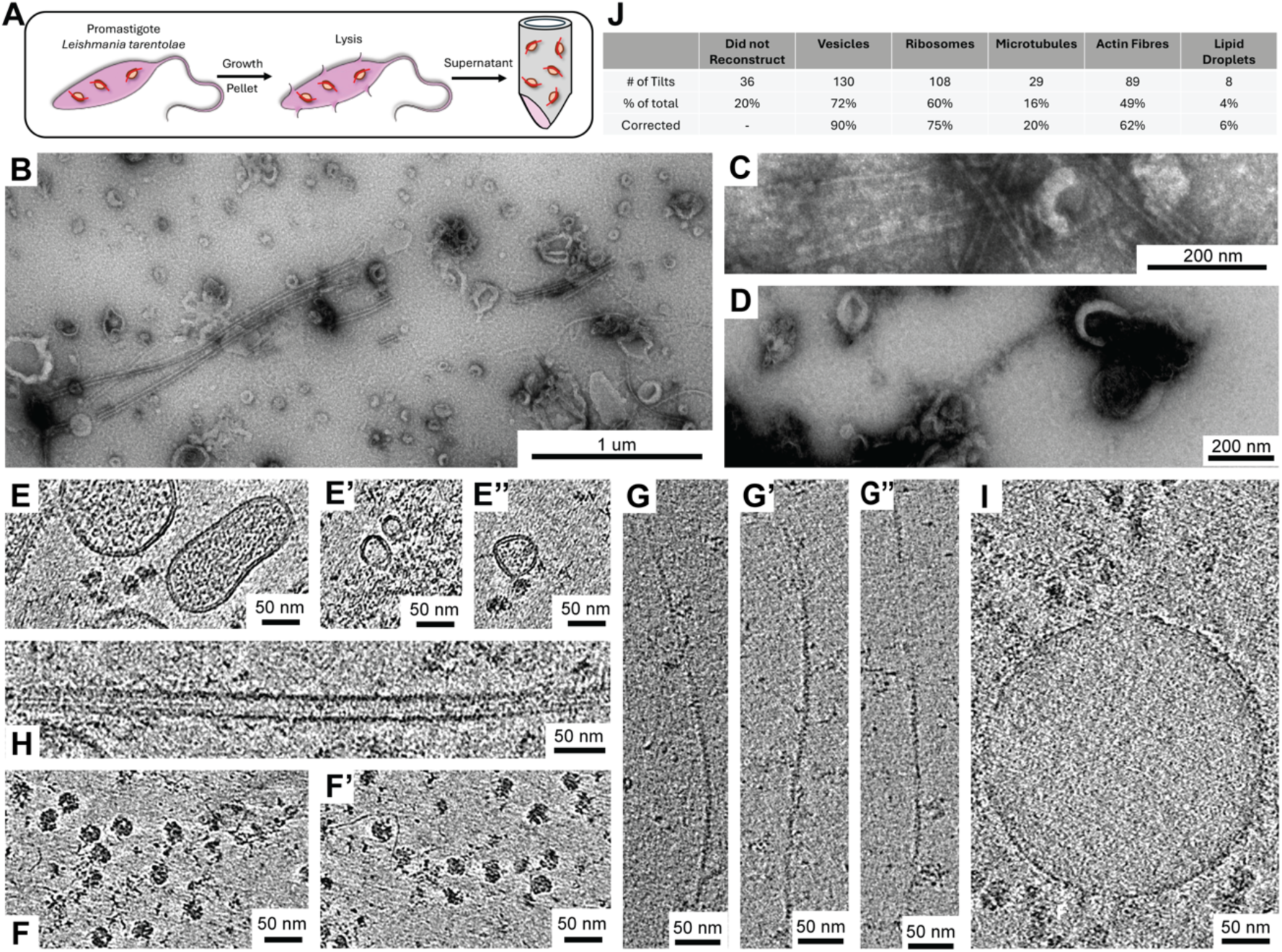
Structural characterisation of the cell-free lysate. (A) Schematic diagram of the cell-free lysate prepared without protein expression. (B-D) Low-resolution negative-stain TEM of native lysate showing abundant microtubules, abundant membrane vesicles and abundant soluble protein adhered to the grid. Cryo-ET screen of frozen lysate showing large and small membrane vesicles (E,E’,E’’), ribosomes (F, F’), actin filaments (G, G’,G’’), microtubules (H) and whole-lipid droplets (I). (J) Scoring of structures present in the native lysate.

### Structure of native *L. tarentolae* microtubules

Given the abundance and structural regularity of *L. tarantolae* microtubules in EM screens screens of LTE, we wanted to first determine that this method of lysate generation and preparation was conducive to high-resolution structural analyses using a well validated target. Therefore, we performed cryo-ET on *L. tarentolae* lysates, with a specific view to solve the structure of native *L. tarentolae* microtubules using sub-tomogram averaging (STA). We developed a Cryo-ET and STA pipeline that employs etomo, Dynamo, crYOLO and Relion5 (Fig. S1) ^26–29^. In agreement with our EM-based screens, abundant and intact microtubules were observed in the lysate (Fig. 3A, B). We quantified the microtubule outer diameter, which measured 29.9 ± 5.6 nm (negative stain) and 27.2 ± 1.8 nm (cryo-ET). These values are slightly larger than the typical microtubules (Fig S3M).

**Figure 3.**
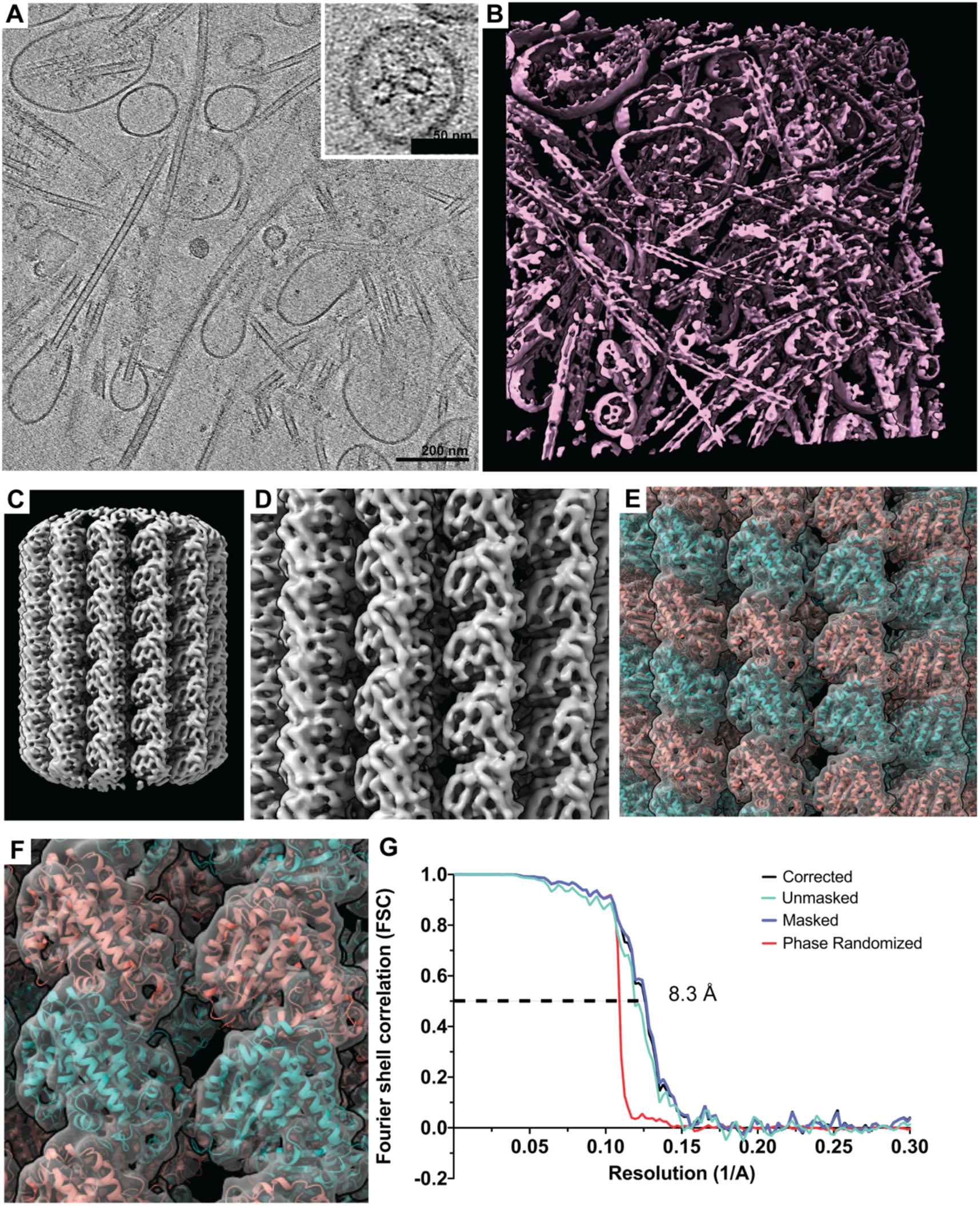
The structure of the *L. tarentolae* microtubules. (A) Slice of a reconstructed tomogram, showing abundant native microtubules. (B) Segmentation of the same volume from (A). No taxol was applied for microtubule stabilsation. (C) Sub-tomogram average of the *L. tarentolae* microtubule, showing secondary structure. (D) Zoomed segmentation of the microtubule reconstruction. (E) Docking of the α- and β-tubulin into the microtubule average. (F) Zoomed-in view, showing the accuracy of the fit of the density map to the predicted structure of *L. tarentolae* tubulin subunits. (G) FSC curve showing 0.5 criterion at 8.3 Å resolution (0.143 FSC criterion at 7.1 Å resolution).

We acquired 208 cryo-electron tomography tilt series at a magnification of 73,000×, with a pixel size of 1.98 Å and a defocus range of 1.0–3.5 µm. Initial processing steps, including motion correction and contrast transfer function (CTF) estimation, were performed on all tilt series using RELION 5.0. Following manual inspection for tomogram reconstruction quality in IMOD 5.1, 42 tilt series were discarded, and the remaining 166 tilt series were selected for further processing. Initial bin4 reconstructions were generated in IMOD 5.1 and used for training microtubule particle picking using crYOLO. Subsequently, particle picking was performed on the 166 bin4 tomograms using crYOLO, yielding 2,354 filaments (44,657 coordinates). These particle coordinates were imported into Dynamo (version 1.1.532) to generate an initial model at bin4 resolution. This model was then used for sequential refinements at bin4, bin2, and high-resolution (bin1) levels, with duplicate particles removed at each step, resulting in 28,971 final coordinates. Tomogram-specific refinements, including CTF refinement and Bayesian polishing, were performed using these 28,971 particles, to produce a final model with a resolution of 8.3 Å (FSC criterion 0.5) (Fig. 3C–G). Notably, this 8.3 Å resolution was achieved using a 200 kV cryo-TEM equipped without an energy filter. The resulting density map resolved detailed secondary structures enabling high-confidence docking of tubulin subunits (Fig. 3; Movie S1). To this end, we first generated AlphaFold 3 structure predictions for the α and β subunits of Leishmania tarentolae and aligned them with the α and β subunit structures of *Homo sapiens* and *Sus scrofa*. The alignments revealed high structural similarity as shown in Fig S1A. Subsequently, we used the Leishmania tarentolae α and β subunit structures for docking into the cryo-electron tomography (cryo-ET) map. The map displayed detailed secondary structures, such as the M-loop, which establishes lateral connections between tubulin heterodimers (Fig. 3F, S2C, and S2D).

Interestingly, although mammalian microtubules are highly sensitive to changes in temperature (a reduction in temperature to 4°C resulting in almost complete depolymerization of the microtubule lattice without taxol stabilisation^30^), the *L. tarentolae* microtubules appear to be particularly resistant to depolymerisation. No taxol was employed in our preparations to stabilize the microtubules. Sample preparation involved temperature changes from 27°C (promastigote culture) to 4°C (lysis), -196°C (snap-freezing), thawing and 4°C incubation prior to plunge-freezing. These changes did not impact microtubule assembly. During its lifecycle, *L. tarentolae* is transmitted between lizards and sandflies and thus subjected to fluctuations in environmental conditions^31^, so it is possible that the stability of the microtubules is a consequence of the need by the parasites to adapt to these changes.

Taken together, our observations demonstrate that the LTE preparation method described here produces samples that are well suited to high-resolution structural analyses. The data also highlight that high-quality, sub-nanometre reconstructions can be generated by sub-tomogram averaging using a 200 kV electron microscope and our optimised data analysis pipeline.

### Correlative cell-free expression and cryo-electron tomography (CC-FLEXCET); application to ASC structural determination

To date, the application of cell-free expression systems to structural biology has been constrained by the limited yield of protein that can be generated using these methods, as well as the reliance on protein purification and protein homogeneity for structural analyses^3^. Having demonstrated that we can generate high quality structural data from cryo-ET of the *L. tarentolae* lysate, we next looked to employ techniques that are routinely applied in *in situ* cryo-ET, to target specific proteins and macromolecular complexes in the lysate^22^. Specifically, we explored coupling cryo-correlative fluorescence microscopy with our cryo-electron tomography methods, to precisely pinpoint targets within the lysate mixture (Fig. 4A).

**Figure 4.**
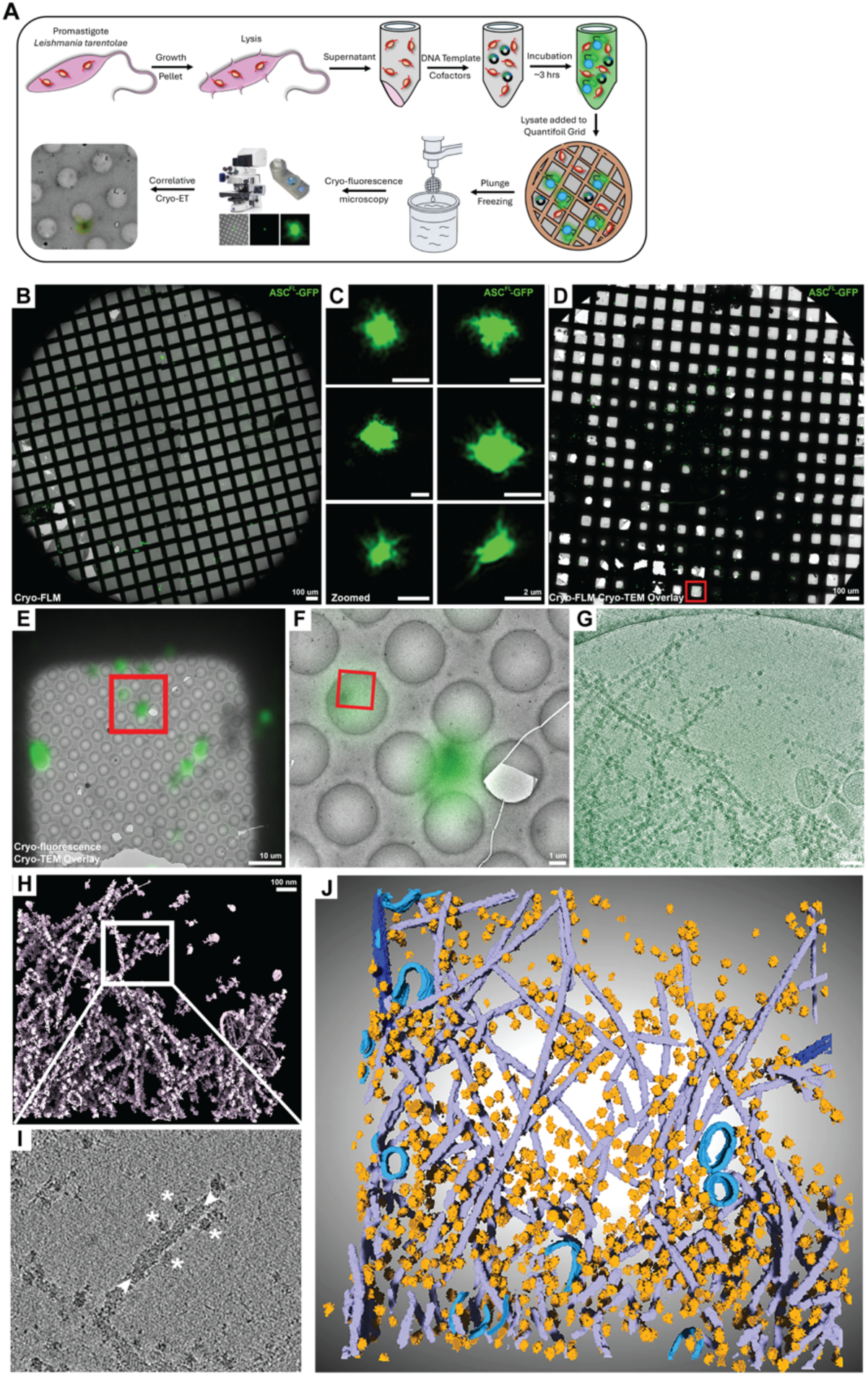
Cryo-Correlative light and electron tomography targeting of expressed GFP-tagged proteins in the *L. tarentolae* lysate and CC-FLEXCET of ASC Specks. (A) Schematic diagram of the CC-FLEXCET workflow. (B) Airyscan cryo-confocal imaging of ASC^FL^-GFP specks in *L. tarentolae* lysate at low magnification. (C) 100X objective imaging of ASC^FL^-GFP specks shows a hairy morphology. (D-G) Overlay of cryo-fluorescence and cryo-EM of lysate expressing ASC^FL^-GFP. The red box denotes the zoomed region of interest. (G) Summed Z-projection for all slices from a reconstructed tomogram showing high GFP intensity correlating with regions of abundant ASC filaments. (H) Segmentation of the whole volume from (G), highlighting the loose meshwork of ASC filaments. The white box shows the area with a single filament decorated by ribosomes. (I) Raw data from (H) showing ribosomes associated with the filament (white asterisk) and a resolvable central pore to the ASC^FL^-GFP filament (white arrowheads). (J) 3D volume segmentation of a tomogram from the ASC dataset, using Amira-Avizo software, displaying ribosomes (orange), ASC filaments (purple), and vesicles (light blue).

As a proof of principle, we performed structural analysis of Apoptosis-associated speck like protein containing CARD (ASC), an adaptor protein essential for the formation of the inflammasome. ASC oligomerisation is critical to stimulate pyroptotic cell death in response to a challenge stimulus. The inflammasome is activated in cells through sensing danger signals, such as LPS or nigericin ^32^, which leads to a structural reorganisation of the NLRP3 protein into an activated state ^33,34^. Upon NLRP3’s activation, ASC is recruited and associates with the NLRP3 complex and oligomerises ^34^. ASC is a 22 kDa protein that comprises an N-terminal pyrin domain (PYD) and C-terminal caspase recruitment domain (CARD), with each domain organised into 6 α-helices linked by a disordered region between amino acids 91 to 115 ^35,36^. ASC filamentous assemblies have been observed *in vitro* by cryo-EM and *in situ* by cryo-ET ^37–39^. *In vitro,* the PYD oligomerises into a filamentous helical arrangement of stacked monomers with an outer diameter of 90 Å and a central pore diameter of 20 Å ^38^. However, it is currently unknown how the CARD is organised around the PYD filament as structural pursuits of full-length ASC (ASC^FL^) have been hindered by the inherent flexibility of the complex^38–40^. We have previously shown that the cell-free expression of ASC in LTE generates large, oligomeric complexes that recapitulate the properties of ASC^FL^ specks in cells^41^. We therefore explored whether the CC-FLEXCET approach could be used to generate the first structural characterisation of the ASC^FL^ filament.

We next sought to characterise the structures of the ASC^FL^ and ASC^PYD^ assemblies resulting from *L. tarentolae* expression. We performed negative-stain transmission electron microscopy on lysates expressing ASC^FL^-GFP and untagged ASC^PYD^ (ASC^PYD^-untagged). We observed that ASC^PYD^-untagged formed abundant filaments in a loose meshwork which were not observed in our lysate-alone controls (Fig. S3A-D). ASC^FL^-GFP also formed filaments, but the organisation of the filaments was different from ASC^PYD^-untagged (Fig. S3E-F). ASC^FL^-GFP adopted larger yet more condensed structures, compared to ASC^PYD^. These complexes accumulated large amounts of uranyl acetate stain and structurally resembled ASC speck formation in cells ^42^. Identifying the specks using negative-stain EM proved challenging, as they had low abundance, and, at low magnification presented as large electron dense patches of accumulated uranyl acetate (Fig. S3E). Only upon high magnification imaging was it possible to resolve the filaments extending and emanating from this dense mass (Fig. S3F-I). To confirm that these specks were comprised of ASC^FL^-GFP, we employed immuno-electron microscopy with a rabbit α-GFP antibody directly conjugated to a 5 nm gold particle. We visualised abundant periodic labelling decorating the ASC^FL^-GFP filaments (more examples of labelled ASC^FL^-GFP specks are included in Fig. S3J-L; Fig. S4A-H). We did not observe immuno-gold labelling of either endogenous microtubules or vesicles present in the lysate (Fig. S4I). These data taken together suggest that LTE expression of ASC^FL^-GFP results in a filamentous assembly of ASC molecules that adopt a regular organisation different from the PYD alone. The abundance of the gold labelling suggests the filament is assembled with the C-terminal GFP-tag on the outside of the fibre.

Our negative-stain EM analyses showed that ASC specks were challenging to find, due to low abundance. This rarity suggests that single particle analysis cryo-EM, which requires large amounts of protein uniformly distributed over the cryo-EM grid, would not be a feasible approach for structure determination of these filamentous assemblies. We therefore employed CC-FLEXCET on the ASC^FL^-GFP specks from the cell-free lysate. We first characterised the morphology of ASC^FL^-GFP specks, by cryo-confocal microscopy on plunge-frozen lysates. Using this method, we observed a low number of bright GFP puncta on each grid that resembled ASC speck observations in cells (Fig. 4B – C^42^). The specks displayed a “hairy” appearance with a diffraction limited core and GFP-rich protrusions extending from the central mass (Fig. 4C). The structures observed by fluorescence imaging also supported our negative-stain and immuno-EM analyses (Fig. S3, S4). We performed cryo-fluorescence imaging, re-registered the GFP-rich complexes in cryo-TEM images (4D-F), and performed guided cryo-ET at the sites of ASC speck formation (Fig. 4G-I). The overlayed data demonstrated that the regions in the centre of the speck were too thick for cryo-tilt series acquisition, but the areas surrounding the central region were ideal and filled with GFP-positive protrusions. Cryo-tomograms of these regions confirmed the presence of extensive filaments, which measured 21.4 ± 1.6 nm. This thickness aligns with our observed filament diameter from negative-stain and immuno-EM analyses (Fig. S3, S4) and *in situ* ASC filaments observed previously ^37^. Intriguingly, we often observed an association between the ASC filaments and ribosomes, although the reason for this is currently unclear (Fig. 4G-J (white asterisk); Movie S2). The cryo-ET data was of high quality, as we could resolve the central pore within the pyrin domain (Fig. 4I; denoted by the white arrowheads) in the raw tomograms. We next sought to solve the structure of the ASC^FL^-GFP from the *L. tarentolae* lysate.

### Sub-tomogram structural analyses of ASC^FL^-GFP filaments

Previous structural studies on recombinantly purified ASC^PYD^ demonstrated that ASC^PYD^ adopted C3 symmetry, featuring a helix with a 90 Å diameter and central pore of 20 Å^38^. Based on the high-quality filaments observed in the cryo-electron tomography (cryo-ET) data, we performed sub-tomogram averaging using the data processing pipeline we developed for *L. tarentolae* microtubules (Fig. 3, Fig. S2). We acquired a dataset of 52 cryo-tomograms of ASC^FL^-GFP filaments at a magnification of 45,000× with a pixel size of 3.19 Å. From these, we selected the nine best reconstructions and used crYOLO for automated particle picking, resulting in 8,476 particles (Fig. S7). This dataset was used to generate an initial model, which revealed the expected organization of the PYD core. Subsequently, we performed 3D classification into three classes and selected class 1 (4,724 particles, 58.3%) for final refinements. These refinements, conducted in RELION 5.0, converged on a model of the ASC^FL^-GFP filament structure using C3 symmetry and local helical symmetry searches with a helical twist of 52.99° and a rise of 13.99 Å (Fig. 5A). The resulting density map clearly resolved the PYD, revealing a central pore with a diameter of approximately 20 Å (Fig. 5B; Movie S3), consistent with previous studies^38,39,43^. Regular density corresponding to the CARDs and GFP tags was also observed around the PYD core as shown in schematic representation (Fig. 5C); however, the overall resolution of ∼20 Å (FSC criterion 0.5) was insufficient to accurately model these domains (Fig. 5A and 5B). We also performed independent reconstructions of ASC^FL^-GFP using either C1 symmetry, C1 symmetry with helical reconstruction, or C3 symmetry with helical reconstruction (Fig. S5A – 5D). All reconstructions showed a similar overall organization of the PYD, with unresolved CARD density and partially resolved outer GFP density in helical reconstructions surrounding the PYD core, as represented in the superimposed map of all reconstructions (Fig. S5C; Movie S3), similar to observations for GFP-AIM2^PYD^ ^44^. The application of helical symmetry improved the overall map quality. This model indicates that, while the organization of ASC filaments is morphologically consistent, significant molecular variability arises from the flexibility observed in the tomograms, filament branching and/or the proximity of ribosomes to the filaments, as discussed below.

**Figure 5.**
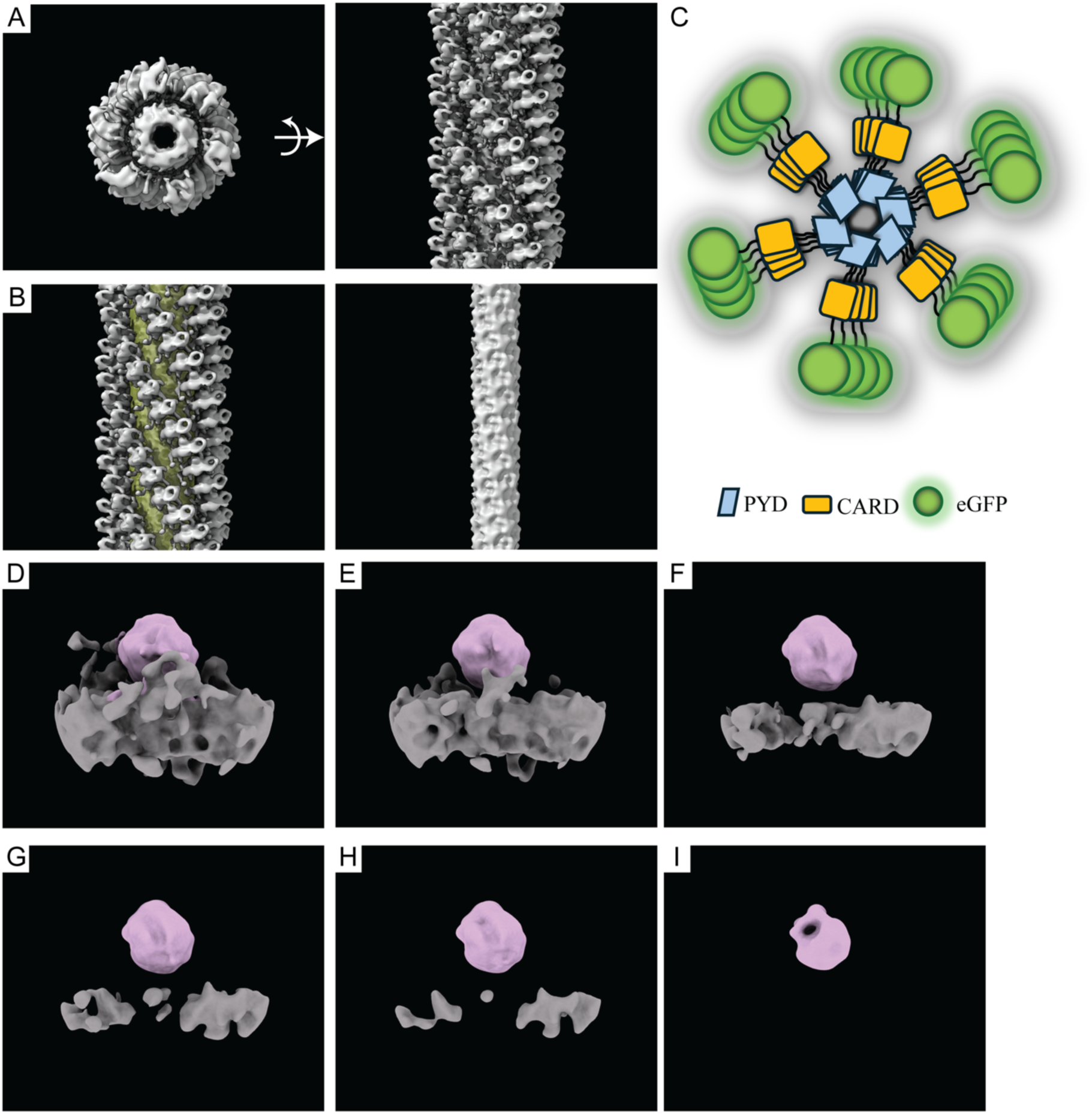
Structure of the ASC^FL^-GFP filament. (A) Sub-tomogram average of ASC^FL^-GFP after 3D classification using (class1) (Fig. S7), showing the overall organisation of PYDs and CARDs. (B) Application of PYD mask and masked average showing PYD core. (C) Schematic representation of PYD – CARD – eGFP assembly showing helical organisation observed in sub-tomogram average structures. (D-I) Different views (thresholded) of refined sub-tomogram average of 97 manually picked particles containing ribosomes (pink) – ASC^FL^ (grey) interactions, showing that the ribosome large subunit faces the filament.

Previous work has shown a specific interaction between ASC filaments and endogenous ribosomes in *in situ* cryo-tomograms of mouse macrophages^37^. We also observed a consistent association between ASC^FL^ filaments and native ribosomes present in the *L. tarentolae* lysate (Fig. 4G–J). Accordingly, we next explored the structure of this association. We initially performed sub-tomogram averaging on the *L. tarentolae* ribosomes, using these same datasets. With 12,310 particles, we solved the native ribosome structure to a resolution of 18 Å (FSC criterion 0.5; Fig. S6A – B). Although the resolution was low, it was sufficient to clearly distinguish between the large and small subunits of the ribosome. In our initial 3D classification of particles from nine tomograms, we did not observe any classes (classes 2 or 3) that revealed specific ribosome attachment to the filaments. Therefore, we manually selected three tomograms based on overall reconstruction quality, ice thickness, filament density, and the presence of ribosome-filament interactions. We then performed 3D classification to sort the ASC filaments into six classes of ASC^FL^-GFP (Fig. S7). In class 3 (Fig. S7), we resolved an association between a single ribosome and the ASC filament, which contained only 789 particles from two of the three tomograms. We attempted to refine this class but could not resolve the ribosome orientation; thus, we manually inspected the picked particles and removed those in dense regions or with ASC^FL^ filaments on both sides of the ribosomes. We focused on sparse regions of the filament and manually picked 97 particles from two tomograms. Refinement of these 97 particles (bin2) is organised such the large subunit is juxtaposed to the ASC^FL^ filament (Fig. 5D-I; Movie. S4).

Sub-tomogram averaging of ribosomes associated with ASC filaments produced a low resolution average that represents the dominant features of ribosome–filament interactions but does not capture potential heterogeneity in individual ribosome orientations. Despite careful manual curation of ribosomes included in the sub-tomogram average dataset, visual inspection of tomograms suggested that ribosomes proximal to the ASC filament may adopt multiple relative orientations. To assess this directly, we performed template matching using our own implementation GAPSTOP^TM^ to determine the orientation of individual ribosomes in situ.

A ribosome reference was generated from the sub-tomogram average of the *Leishmania tarentolae* ribosome (Fig. 6A), with labels marking the large (LSU) and small (SSU) ribosomal subunits. Initial attempts at tomogram-wide template matching using a full grid search were unsuccessful due to strong thickness gradients across the tomogram and interference from dense filamentous structures, resulting in the detection of low-quality matches distributed throughout the volume. Because our aim was to resolve ribosome orientations specifically in the vicinity of ASC filaments, we instead manually curated 218 ribosome positions using Napari and extracted corresponding sub-volumes for targeted template matching.

**Figure 6.**
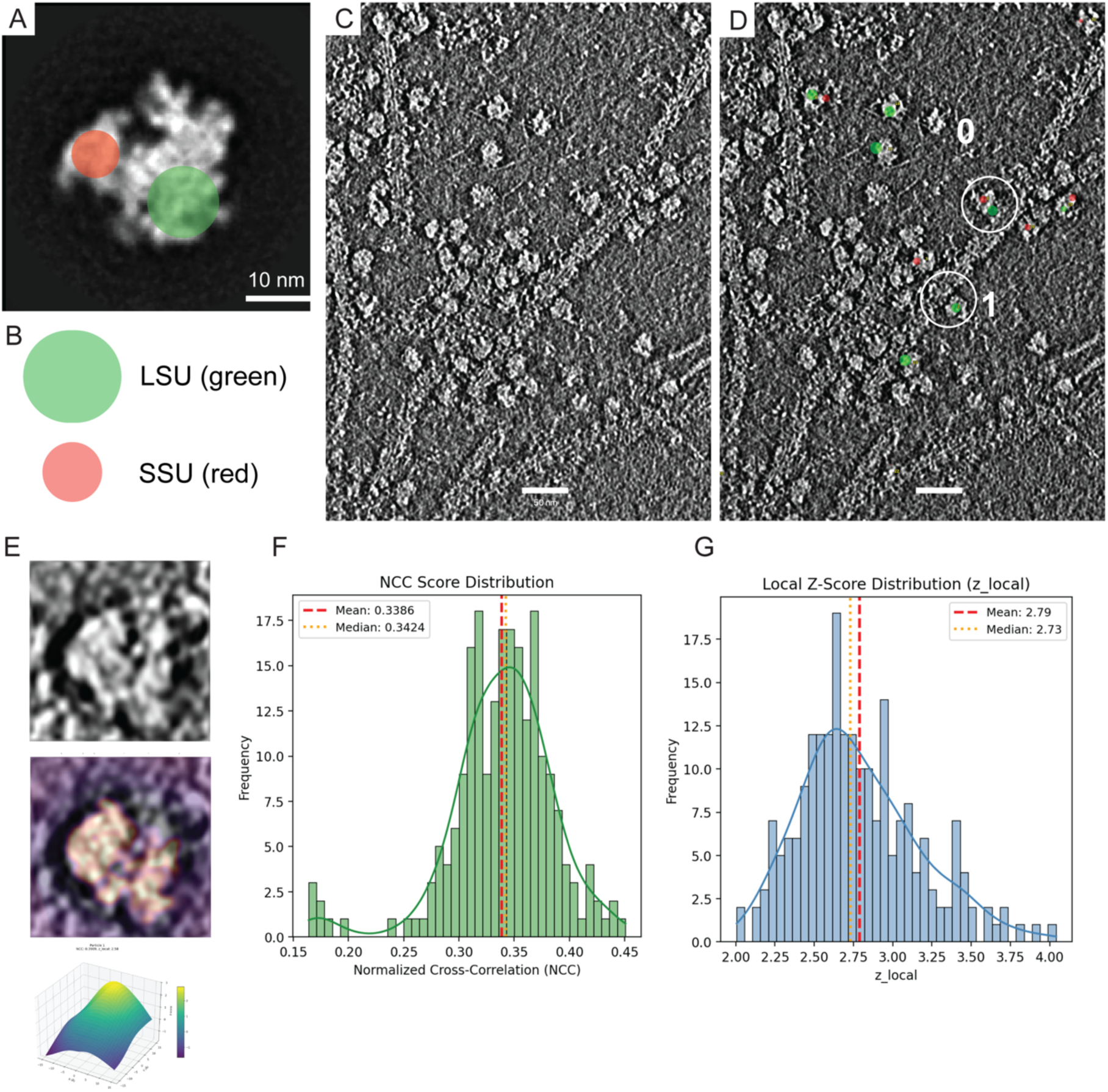
Template matching reveals heterogenous ribosome association with ASC filament. (A) Ribosome template derived from sub-tomogram averaging of *L. tarentolae* ribosomes. Positions of the large ribosomal subunit (LSU, green) and small ribosomal subunit (SSU, red) are indicated and used as orientation markers for subsequent analyses. Scale bar, 10 nm. (B) Schematic representation of LSU (green) and SSU (red) marker labels used to classify ribosome orientation relative to the ASC filament. (C) Representative tomographic slice showing an ASC filament with the dense and heterogeneous mixture of ribosomes are embedded. Scale bar, 50 nm. (D) Same region as in (C) with projected LSU (green) and SSU (red) markers overlaid following template matching and local alignment. Circles highlight representative ribosome exhibiting distinct orientations relative to the filament surface. (E) Representative ribosome subvolume before (top) and after (bottom) template alignment overlay, and Local z-score surface for a representative ribosome, demonstrating a distinct correlation maximum. **(F)** Distribution of normalized cross-correlation (NCC) scores obtained from template matching across all curated ribosomes. The distribution (mean and median indicated) reflects consistent template alignment quality within the dataset. (G) Distribution of local z-scores (z_local) computed from spatially local cross-correlation backgrounds surrounding each particle.

For each curated ribosome sub-volume, the template was locally aligned to the sub-volume, and the resulting pose parameters (Euler angles and translational offsets) were used to back-project LSU and SSU marker labels into the original tomogram. This allowed direct visualization of ribosome orientation relative to the ASC filament (Fig. 6C–D). Matches with normalized cross-correlation (NCC) scores below 0.2 were excluded from further analysis. The remaining particles showed a unimodal NCC score distribution with a mean of approximately 0.30 (Fig. 6F), indicating consistent template alignment across the curated dataset. Local z-scores, computed from the spatial distribution of cross-correlation values surrounding each particle, showed a broad but well-defined distribution (Fig. 6G), supporting the statistical significance of individual matches. The magnitude of these z-scores is lower than values reported in recent tomogram-wide grid-search–based template matching approaches, reflecting our targeted matching strategy in which candidate positions were manually curated and background statistics were estimated locally rather than pooled across the entire tomogram. Consequently, z-scores in this study should be interpreted as conservative, position-specific measures of alignment significance rather than global detection scores.

Analysis of ribosome orientations revealed substantial heterogeneity in how ribosomes associate with the ASC filament. Among the 218 curated particles, 126 ribosomes (57%) were oriented with the large ribosomal subunit juxtaposed to the filament surface, while 92 ribosomes (43%) displayed either the small subunit facing the filament or their relative orientation could not be unambiguously assigned due to limited contrast or partial occlusion by surrounding density. Representative example of distinct ribosome orientations are shown in Fig. 6D.

Taken together, these data highlight that full-length ASC is promiscuous in its oligomerisation and adopts flexible arrangements, which inhibit solving the structure of the complex at sub-nanometre resolution. Nonetheless, we observe an overall consistent organisation of the ASC filament that possesses a more rigid helical arrangement of the PYD core with C3 symmetry and a peripheral network of CARDs that are flexibly oriented around the PYD core in a twisted and stacked organisation. In this context, ASC filaments can form an association with the ribosome with a slight preference for positioning of the large ribosomal subunit in closer proximity to the filament surface. While this association is apparent at the ensemble level when observed by sub-tomogram averaging technique, individual particle level orientation analysis reveals that ribosomes proximal to ASC filament adopt multiple orientations with more than 50% particles presenting their large subunit to filament but alternative orientations are also widely represented. These observations indicate that ribosome – ASC filament interactions are structurally heterogenous and that this heterogeneity is not captured by sub-tomogram averaging alone. Finally, and critically, we demonstrate that CC-FLEXCET system can be utilised to perform structural characterisation on proteins of unknown structure rapidly and without biochemical purification.

## Discussion

Recombinant systems are often tailored for each protein; some proteins express and fold well with traditional *E. coli*-based methods, while other proteins can require a slow and tedious cycle of optimisation and validation. This often involves switching between different expression systems and time-consuming optimisation of purification conditions, aiming to generate higher quality protein or protein complex formation for structural analyses. Here we have developed and characterised CC-FLEXCET, a simplified cell-free system that can be applied to structural studies of proteins without the need for isolation and purification. We show that although *L. tarentolae* lysates contain endogenous proteins and membrane structures observable by cryo-EM, these contaminants do not prevent the direct structure determination of exogenously expressed proteins by cryo-ET. Enabled by fluorescent tagging of proteins of interest we can readily target macromolecular complexes with cryogenic-confocal microscopy coupled with cryo-correlative electron tomography. We utilise this approach to express ASC^FL^-GFP and demonstrate that cell-free expression recapitulates the structure of ASC specks from cells. We link the above targeting approaches with sub-tomogram averaging to perform the first structural analyses of full-length human ASC filament without the requirement for biochemical purification. The cell-free expression lysate utilised here is derived from *L. tarentolae,* but it is important to note that other cell-free systems have been used for structural studies^11,12,14,15^ and the pipeline presented here is readily applicable to those other systems. The method therefore has the potential to be widely adopted as a tool for cell and structural biologists looking to analyse protein structures in more complex systems, without requiring the complete complement of equipment for, and complexities that come with, the full *in situ* cryo-ET workflow.

The characterisation of the ASC full-length structure has previously been complicated by the difficulty of purification and its propensity to constitutively oligomerise. Here we have circumvented these problems by studying ASC filaments directly in the LTE cell-free system. It was initially proposed that ASC assembles into hollow donut-like arrangements in the cytoplasm in response to inflammasome activation^45,46^. Later it was shown that ASC adopts a dense nucleated core of protein within the speck and from this core filamentous ASC polymers extend into the cytoplasm^38,42^. Here we show that the gross morphology of the ASC speck structure formed via cell-free expression shows a morphology consistent with later cellular observations^38,42^. Our cryo-fluorescence microscopy (Fig. 5A – B) and our lower-resolution conventional transmission electron microscopy analyses highlight the propensity for ASC to form a dense central domain with filaments extending from this core (Fig. 5; Fig. S3E, J; Fig. S4). Using the CC-FLEXCET protocol for cryo-electron tomography, we could not only observe the formation of a defined ASC filament (Fig. 6F-I), but we could also resolve interactions and organisations that have previously only been observed *in situ*. In immortalized mouse bone marrow derived macrophages, Liu et al (2023)^37^ observed that ASC formed a loose meshwork of fibres in the ASC speck, and that ASC filaments were enriched in, and associated with, ribosomes *in situ.* We observed a similar loose arrangement of ASC filaments as well as a frequent association between the *L. tarentolae* ribosome and ASC filaments (Fig. 6D-I). Lu et al (2023)^37^ proposed that this ribosome:ASC interaction represents a method for cells to control signalling or rapid response to danger signals. However, it is unlikely that this ribosomal association represents newly synthesised ASC in cells, as the specked form arises rapidly after inflammasome stimulation. In our cell-free system, the frequent yet irregular association between ASC filaments and ribosomes suggests that this interaction is biologically relevant but does not preclude the possibility that this binding is a consequence of real-time translation of the ASC protein with concurrent assembly of the filament or, potentially, stalled translation. We used template matching analyses of the ribosome:ASC association to show a slight preference for the large subunit in association with the ASC filament. Ribosomes are often observed relocating to other structures as a reaction to cellular stress, and it is possible that this association represents an analogous response^47^. Regardless, it remains unclear as to what the role this association is playing in inflammasome signalling.

Here, we generated the first reconstruction of the filamentous form of the full-length human ASC protein. While our structural pursuits could not generate a sub-nanometre resolution reconstruction of the ASC filament, we could still derive valuable structural information from the analyses that we performed. We identified that the full-length ASC assembles into a filament with a PYD core, a central pore and peripheral CARDs. The arrangement of the core PYD filament was more rigid and estimated at ∼20 Å resolution (FSC criterion 0.5). The apparent flexibility of the CARDs, however, makes it more challenging to model. Our reconstructions show a regular stacked arrangement of 6 densities twisted around the outside of the filament, which presumably correspond to the summed density of both the CARD and the GFP-tag. While our structural model has had both C3 and helical symmetry applied, the retention of this density indicates this stacked arrangement is at least somewhat regular but, unfortunately, the separation of these densities into two different domains (CARD and GFP) was not possible at our attained resolution. It is likely that the ring like domain observed in this reconstruction corresponds to the β-barrel of GFP.

Homotypic CARD:CARD interactions have been shown to be critical in numerous mutational studies, highlighting that the C-terminal domain is essential for both speck and filament formation ^42,48^. Moreover, CARDs alone have been shown to form higher-order filament-like structures by cryo-EM through CARD interaction interfaces (type I, type II, type III) ^49^. The stacked association of CARD/GFP domains in our model suggests that the CARD undergoes some homotypic association through these vertical interaction interfaces, but it is unclear from these averages how this kind of association of CARDs could be possible in the context of the full-length ASC filament^49^ without resolvable lateral interactions. Despite this, the spacing of these CARDs would leave adequate distance to facilitate binding of, and interaction with, pro-caspase-1 CARDs for subsequent downstream inflammasome signalling. It is important to note that the lower-resolution reconstruction that we generated for ASC was not a consequence of the data quality, the microscope, the data processing, nor the sample preparation as we demonstrated that we could use this same approach to generate an 8.3 Å resolution reconstruction of the *L. tarentolae* microtubules. Rather, we believe this is a consequence of the inherent flexibility of the ASC filament which has hindered higher-resolution analyses of the full-length protein for over a decade.

The CC-FLEXCET pipeline offers a rapid pathway from protein expression to data acquisition within a single day and is particularly well suited to studying the structures of proteins that form large macromolecular assemblies such as ASC. In its full-length form, ASC does not distribute uniformly on grids, and as such is not conducive with single particle analyses - the behaviour of this protein is such that its imaging requires targeting using correlative microscopy and sub-tomogram averaging. However, this does also highlight a caveat to the method; as we are targeting complexes by fluorescence microscopy, a sufficient quantum of fluorescence that can be resolved by cryo-fluorescence microscopy is required for correlative cryo-EM. While we are performing confocal microscopy with an Airyscan detector, we are not in the range of single molecule detection. Thus, it is unlikely that without higher order oligomerisation to generate sufficient clustering of fluorophores, CC-FLEXCET would be able to generate distinct fluorescence signals to distinguish between monomeric and small molecular weight complexes. We believe, in its current form, the system is limited to studies of larger protein assemblies but alternative approaches could yield broader application. By tethering smaller complexes to other structures present in the lysate (such as microtubules, membranes, or lipid droplets) or through functionalised affinity cryo-EM grids ^50^ we hypothesize we could circumvent this limitation. This caveat is also balanced with some significant advantages. This cell-free system has been used extensively for characterising protein-protein interactions by single molecule coincidence detection ^8,10,41^. Here, we are not limited by a single fluorophore and can use this expression approach to target specific protein complexes, based on the ratio of GFP to other expressed protein with a different fluorophore like mCherry. As an extension of this, our fluorescence detection can happen without biochemical isolation and as such, we could potentially explore this approach to look at fragile complexes that are unable to be isolated from cells without extensive crosslinking, such as the Commander complex^51^.

The development of methods to analyse complicated samples, such as those required for *in situ* cryo-ET, are facilitating a change in the way we can approach solving protein structures^17^. The reductionist structural approach to analysing a homogenous protein sample in a small number of conformational states, while informative, is not sufficient on its own to understand complex protein assembly and behaviour in the cell. Conversely, a purely cell-biological approach to understanding protein function and structure, by performing true *in situ* molecular sociology, is currently impeded by the complexities of the cell, the preparation of samples for structural analyses and often the relatively low abundance of the proteins of interest. Here, we have described a pipeline that operates in the middle ground between these two approaches; this system can provide information beyond the structure of an individual purified protein or protein complex yet represents a less complex mixture than the native cellular environment. To date, methods that occupy this space have largely been based on crude lysates extracted from cells of interest, for structural analysis on native proteins ^52–54^. Alternatively, cell-free approaches have been applied to solve structures of several proteins by single particle cryo-EM, but only after performing protein purification^11,12,14,15^. Our CC-FLEXCET method provides several advantages. Using CC-FLEXCET, we can express our desired proteins of interest within a lysate and apply targeting methods developed for *in situ* cryo-ET to re-register our proteins of interest within the lysate. This cell-free approach provides a unique opportunity to understand protein structure as well as image protein interactions within a simpler framework than the cellular environment.

## Experimental Procedures

### *Leishmania tarentolae* culture and lysate production

*L. tarentolae* promastigotes were obtained from Jena Bioscience GmbH. Cells were cultivated in a Biostat A fermenter in modified terrific broth with both glycerol and glucose (TBGG) supplemented with 0.05% w/v hemin at 27 °C ^55^. Cells were pelleted for at 2.500 x*g* and the pellet was washed with sucrose extraction buffer (SuEB) containing 250 mM sucrose, 100 mM potassium acetate and 3 mM magnesium acetate. *Leishmania* was resuspended in SuEB, transferred into a cell disruption bomb (Parr Instruments) and incubated on ice under 70 bar nitrogen pressure for 30 minutes ^56^. Release of the pressure stimulated nitrogen cavitation-based lysis of the *L. tarentolae*. The supernatant was subjected to centrifugation at 30,000 x*g* for clarification, prior to the addition of anti-splice DNA leader oligonucleotides (10 μM). The lysate changed to a 45 mM HEPES, pH 7.6, containing 100 mM potassium acetate and 3 mM magnesium acetate buffer, using illustra NAP-25 columns, snap-frozen in liquid nitrogen and stored at −80 °C until required.

### *L. tarentolae* cell-free expression

Plasmids were constructed using the pCellFree backbone ^57^. Constructs were cloned into the following Gateway destination vectors: N-terminal GFP-tagged (pCellFree_G03), N-terminal mCherry-tagged (pCellFree_G05), C-terminal eGFP-tagged (pCellFree_G04) or C-terminal mCherry-cMyc-tagged (pCellFree_G08). The following plasmids were used: pCellFree_G04-ASC (ASC^FL^-GFP), pCellFree_G01-PYD (PYD-untagged). Protein expression was performed as described ^56^, using 20 mM of DNA template in 10 μL to 20 μL of *L. tarentolae* extract lysate, for 3 h at 27 °C, unless stated otherwise. Samples were processed immediately for either fluorescence correlation spectroscopy or plunge-freezing.

### Transmission electron microscopy of *Leishmania tarentolae* cells

Cells were fixed in 5% glutaraldehyde, diluted 1:1 in TBGG to a final concentration of 2.5% glutaraldehyde, for 1 hour at room temperature. Briefly, cells were pelleted after fixation, resuspended in 0.1 M sodium cacodylate buffer, adhered to 35 mm poly-L-lysine coated cell culture dishes, then processed exactly as described previously ^58^. Resin polymerisation, ultrathin sections were cut on a UC6 ultramicrotome and imaged on a JEOL1011 transmission electron microscope at 80 kV.

### Negative stain transmission electron microscopy

200 mesh carbon coated formvar grids were glow-discharged and incubated for 10 minutes with control *L. tarentolae* lysates or lysate expressing proteins of interest. Grids were washed 5 times for 1 minute in phosphate buffered saline (PBS), then water washed 5 times rapidly prior to negative staining with 1% aqueous uranyl acetate. Grids were imaged at 80 kV on a JEOL 1011 transmission electron microscope fitted with a Morada camera.

### Immuno-electron microscopy

Lysates were adhered to grids as described above, then fixed in 4% paraformaldehyde in PBS, washed in PBS 2 times for 5 minutes, quenched in 10 mM glycine, washed again in PBS, and blocked in 0.1% bovine serum albumin and 0.1% fish skin gelatin in PBS for 20 minutes. Grids were incubated with 4.5 nm gold directly conjugated to rabbit anti-GFP antibody for 30 minutes, then washed 5 times for 5 minutes in PBS. Samples were washed 5 times rapidly in water and counterstained with 0.4% uranyl acetate in 2% methylcellulose, then imaged on a JEOL 1011 transmission electron microscope fitted with a Morada camera.

### Plunge freezing

Lysates expressing PYD-untagged, ASC^FL^-GFP were prepared as described above. 200 mesh carbon film Quantifoil copper grids (R3.5/1) were plasma cleaned and 4 uL of lysate was added to each grid and plunge frozen into liquid ethane on a Leica EM GP2. Grids were stored in liquid nitrogen until cryo-confocal imaging.

### Cryo-confocal microscopy

Frozen grids were loaded onto a Linkam CMS196 cryo-stage fitted with an Autofil dewar. The stage was mounted onto a Zeiss LSM900 upright confocal microscope with an Airyscan 2 detector. Low magnification 5X fluorescence imaging was performed to map out grids using the 3-grid holder for the Linkam stage. A digital zoom of 0.45xMag provided an image of the whole grid and these 5X maps (one for each grid) were acquired on a custom holder layout implemented in Zen Blue (Zeiss), using 488nm excitation for GFP with a separate channel for PMT acquisition of transmitted light on all grids. Automated imaging was performed as follows (see Fig. 6C). Higher resolution maps were acquired on a 10X objective with a digital zoom of 2.5X with over-sampling on the Airy detector for all three grids. GFP channel was acquired with 488 nm laser excitation on the Airy detector with a laser power of 1-2%. A transmitted light channel was also acquired using the PMT. This channel is used for correlating between the light microscopy and the cryo-TEM, as each hole on the quantifoil grid is observable in both light and EM and represents an essential fiducial marker for re-registration (Tillu et al 2024). Autofocus at each position was scripted in Zen Blue. For high-resolution fluorescence imaging (Fig. 6B), z-stacks of GFP puncta were acquired on 100X (0.7 NA objective) using batch acquisition and autofocus. Maximum intensity z-projections were generated of the z-stack data.

### Cryo-electron tomography

Frozen grids were loaded into a Glacios2 Cryo-Transmission Electron Microscope (Thermo Fisher Scientific) fitted with a Falcon 4i Direct Electron Detector (Thermo Fisher Scientific) under the control of the Tomography 5 software (Thermo Fisher Scientific). Batch slab-like cryo-tilt series were acquired at 3° increments from -60° to +60° (41 images), with a dose-symmetric acquisition scheme, using multi-position parallelisation. 10-frame movies were acquired at each tilt angle, using electron counting mode. Total dose of each tilt series was ∼90 e^-^/Å^2^, with a pixel size of 1.98 Å for the microtubule data and 3.19Å for the ASC acquisition. Tilt series were batch reconstructed using patch tracking implemented in the eTomo software ^59^. Batch reconstructs were used to screen for data quality.

### Cryo-correlation

Cryo-TEM grid atlases were exported and manually aligned to the 10X fluorescence maps using Adobe Photoshop during acquisition on the cryo-TEM, as described previously (Tillu et al. 2024). Cryo-TEM grid square overviews (700X magnification) were exported and aligned regions of interest identified in the fluorescence maps. Alignment, correlation, and re-registration were performed using the Quantifoil holes as fiducial markers in Photoshop. Higher magnification search maps (3400X) were acquired as tiles in Tomography 5 for final high-accuracy positioning for batch tilt series positioning.

### Sub-tomogram averaging

For each dataset, the pipeline workflow (Fig. S1) was followed. Dose-fractionated tilt series were imported into RELION 5.0^29^, for pre-processing prior to tomogram reconstruction and sub-tomogram averaging. Motion correction was performed using RELION’s implementation of a MotionCor2-like algorithm, and CTF estimation was conducted using the CTFFIND-4.1 implementation in RELION 5.0^29^. These pre-processing steps were applied to all tilt series in each dataset. Subsequently, each tilt series was manually inspected, and images exhibiting ice contamination, grid bars, or significant movement at higher tilts were excluded. The refined tilt series metadata were exported as a STAR file from RELION 5.0.

Refined tilt series were imported into eTomo^59^ (IMOD 5.1.1) for alignment using Batchruntomo. Alignment quality was assessed using the mean residual error (reported in nm by eTomo), and tilt series with a residual error greater than 2.5 nm were excluded. Aligned tilt series were imported back into RELION 5.0 for tomogram reconstruction using the weighted backprojection method. Bin4 tomograms were generated with CTF correction (--ctf flag) and used for two purposes: (1) training models for microtubules, hASC filaments, and ribosomes in crYOLO^27,28^ for automated particle picking, and (2) generating initial bin4 models in Dynamo with C1 symmetry. These initial models were low-pass filtered to 60 Å, resulting in featureless rod-like structures for microtubules and hASC filaments, serving as starting points for bin4 refinement in RELION 5.0.

All three datasets achieved Nyquist resolution at bin4. Sub-tomogram averaging proceeded by extracting particles at bin2, followed by further refinements to generate bin1 references. Duplicate particle coordinates were removed at each refinement stage, based on minimum distance thresholds: 80 Å for microtubules and 40 Å for hASC filaments. Helical reconstruction was performed for the microtubule and hASC filament datasets, with local searches for helical twist and rise conducted within known ranges of helical symmetry parameters. For microtubules, the helical twist search ranged from -24 to -30 degrees, and the helical rise search ranged from 6 to 12 Å, yielding optimal values of -27.68 degrees and 9.95 Å, respectively. For hASC filaments, the helical twist search ranged from 50 to 55 degrees (based on published PYD core helical symmetry), and the helical rise search ranged from 11 to 15 Å, resulting in optimal values of 52.99 degrees and 13.98 Å, respectively.

Tomography-specific refinements (tomo refinements) were performed in RELION 5.0, including CTF defocus refinement and Bayesian polishing with per-particle motion refinement. These refinements were conducted iteratively, followed by auto-refinement in RELION 5.0, to generate the final sub-tomogram average with post-processing. A uniform B-factor sharpening of -100 Å^2^ was applied to all maps during post-processing.

### Template Matching

Template matching on cryo-electron tomograms was performed using a custom GPU-accelerated pipeline built with our own implementation of GAPSTOP^TM^ to detect ribosome orientation proximal to ASC filaments at manually curated ribosome positions while accounting for missing-wedge effects. Template matching was restricted to predefined coordinates rather than performed as an exhaustive search. For each coordinate, a cubic sub-tomogram of fixed size (typically 136³ voxels, bin1 pixel size – 3.19 Å) was extracted and centered on the provided position, with zero-padding applied at tomogram boundaries. Reference templates were resampled to match the tomogram voxel size, then center-cropped or padded to the sub-tomogram box size. An optional soft-edged spherical mask was applied to restrict correlation to central molecular density and suppress edge artifacts. The same mask was applied consistently to both templates and sub-tomograms.

Sub-tomograms and templates underwent identical pre-processing. A Fourier-space bandpass filter with raised-cosine transitions suppressed low-frequency background variations and high-frequency noise using user-defined cut-offs (typically 400 Å high pass, 20 Å low pass). Filtered volumes were normalized to zero mean and unit variance, with statistics computed within the masked region when applicable. Template matching was performed over discrete Euler angles using the ZYZ convention (RELION standard). A coarse angular grid (30° spacing) was evaluated first, followed by hierarchical local refinement around top-scoring orientations using progressively smaller angular steps (15° → 7° → 3° → 1°) until convergence. Template rotations were implemented using GPU-accelerated affine transformations and evaluated in batches for computational efficiency. To compensate for anisotropic resolution from limited tilt acquisition, a missing-wedge mask was constructed in Fourier space based on the tilt range (±60°) and tilt axis orientation. During correlation, Fourier transforms of both sub-tomogram and template were multiplied by this mask such that only mutually sampled coefficients contributed to the score. Normalized cross-correlation (NCC) was computed in Fourier space, and correlation volumes were shifted such that zero displacement corresponded to the central voxel. Correlation peaks were identified within a restricted search window (±15 voxels) centred on zero shift, with subpixel refinement performed by parabolic interpolation along each axis. For each particle, a local z-score (z_local) was computed to assess peak significance by estimating background statistics from correlation values within the search window, excluding a small spherical region (radius = 2 voxels) around the detected peak.

## Acknowledgements

NA is supported by a Human Frontier Science Program (HFSP) Grant (RGP/011/2023) and an Australian Research Council (ARC) Discovery Project (DP260101196). RGP was supported by an ARC Laureate Fellowship (FL210100107) and Discovery Project (DP260101196). This research is part of a project that has received funding from the European Research Council (ERC) under the European Union’s Seventh Framework Programme (FP7-2007-2013) (Grant agreement No. 101071784). BK was supported by NHMRC Investigator funding (2025931). This work is supported by funds to BMC from the National Health and Medical Research Council (NHMRC) Investigator Grant (APP2016410). The authors acknowledge the use of the Microscopy Australia Research Facility at the University of Queensland.

## Author Contributions

Initial Concept: NA, RGP, YG

Concept development: NA, VT

Technical assistance: JS, DH, YG, JR, ON, ES

Structure determination and analysis: NA, VT, KC, BMC

CryoET studies: VT, NA, JR

Data analysis and figure generation: VT, NA

Funding acquisition: NA, YG, RGP, BMC, BK

Supervision: NA

Writing – 1st draft: NA, VT

Writing – review & editing: NA, VT, RGP, BMC, BK

## Data availability

All structures to be uploaded to the EMDB upon acceptance of the manuscript.

## Supplemental Movies

Movie S1 – 3D rendering of the sub-tomogram average map of a microtubule filament with docking of α- and β-tubulin subunits.

Movie S2 – 3D cryotomographic reconstruction of a tilt series showing ASC^FL^-GFP filament with ASC-ribosome interactions.

Movie S3 – 3D rendering of sub-tomogram averages of ASC^FL^-GFP with superimposition of C1 (grey), C1 helical (yellow), and C3 helical (cyan) reconstructions.

Movie S4 – Two-color 3D rendering of the sub-tomogram average of ASC (grey) and ribosome (pink) interactions with different thresholds.

**Supplementary Table 1.** Data collection and statistics.

| Data collection |  |  |  |
| --- | --- | --- | --- |
| Data collection | Leishmania Microtubule | Human ASC<sup>FL</sup> | Leishmania 80S Ribosome |
| Transmission EM | Glacios 2 | Glacios 2 | Glacios 2 |
| Magnification (nominal) | 73,000 | 45,000 | 45,000 |
| Accelerating voltage (kV) | 200 kV | 200 kV | 200 kV |
| Total electron dose (e<sup>-</sup> per Å<sup>2</sup>) |  |  |  |
| Defocus range (µm) | 1.0 µm – 3.5 µm | 4 µm | 4 µm |
| Tilt series recorded/used for processing | 208 / 166 | 52 / 9 | 52/52 |
| Pixel size (Å) | 1.98 Å (native)<br>0.99 Å (8k) | 3.19 Å | 3.19 Å |
| Data processing and map reconstruction |  |  |  |
| Data processing mode (native or eer upsampling) | EER (8k * 8k) | Native (4k * 4k) | Native (4k * 4k) |
| Image-processing package | Relion 5.0, Dynamo | Relion 5.0, Dynamo | Relion 5.0, Dynamo |
| Initial picked particles or subtomograms | 44657 | 8746 | 12310 |
| Final used particles or subtomograms | 28971 | 4724 | 3721 |
| Map resolution | 8.3 Å | 20 Å | 18 Å |
| FSC threshold | 0.5 | 0.5 | 0.5 |
| Map sharpening B factor (Å<sup>2</sup>) | -100 | -100 | -100 |

## Supplementary Figures

**Figure S1.**
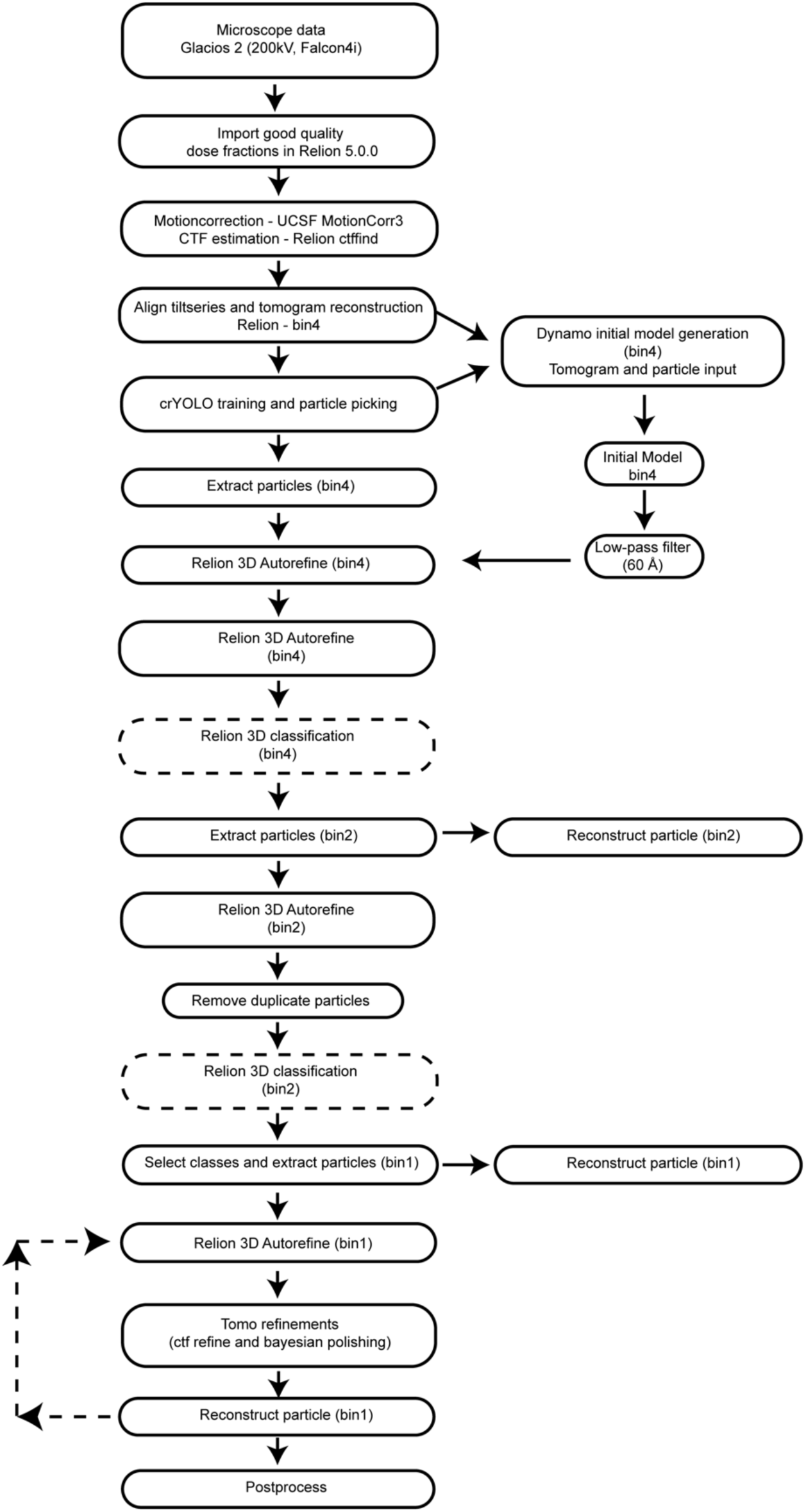
Data processing pipeline for sub-tomogram averaging. A sample template of data processing pipeline used to generate subtomogram average structures using Relion 5.0, Dynamo and IMOD.

**Figure S2.**
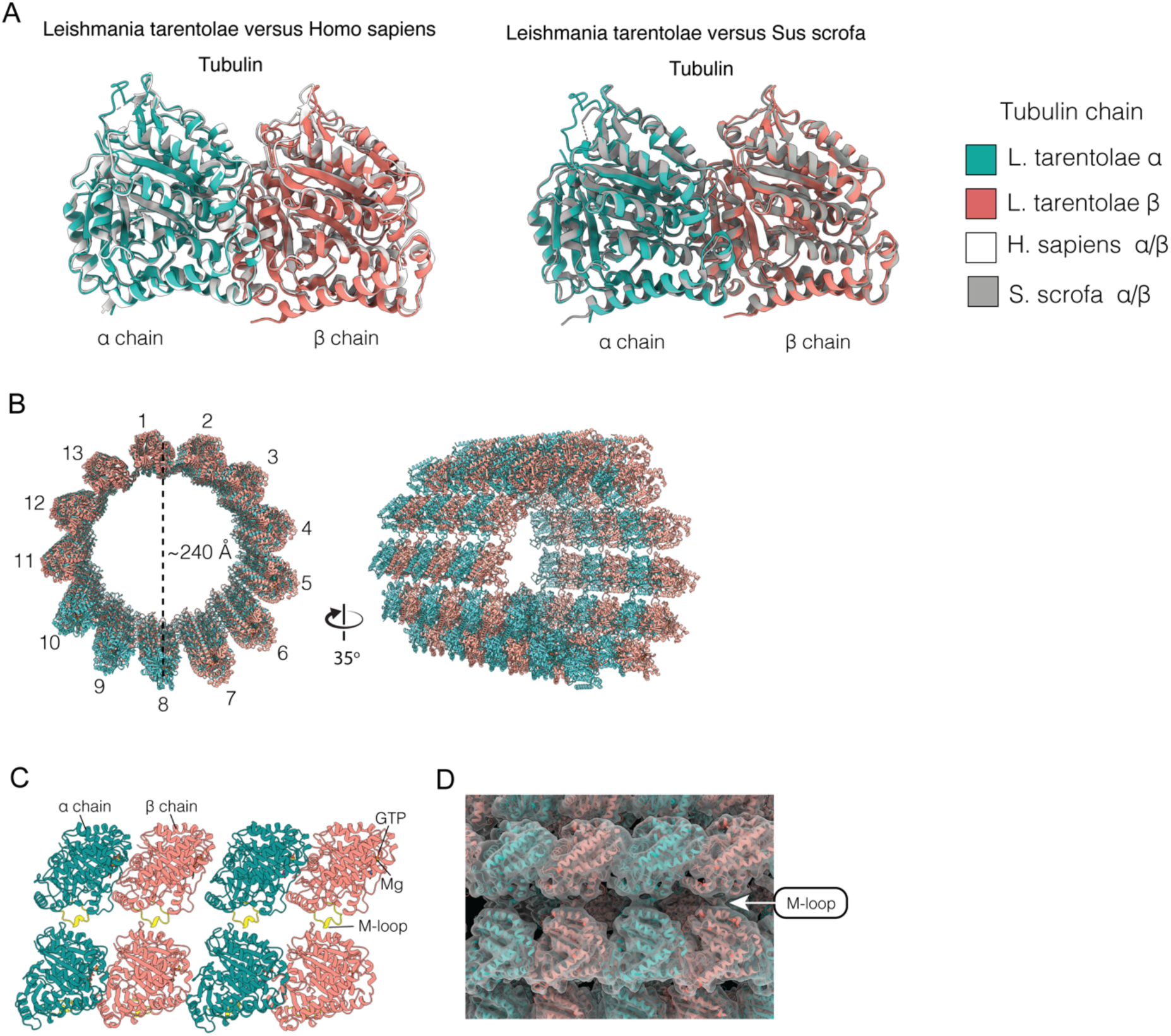
L. tarentolae tubulins form canonical microtubule filaments. (A) Structural superimposition showing that the AF2-predicted L. tarentolae αβ-tubulin dimer model is highly similar to the H. sapiens (PDB ID: 5JCO) and Sus scrofa structures (PDB ID:3J6E). (B) Docking of the L. tarentolae microtubule model into the density map generated from STA. The L. tarentolae microtubule exhibits a canonical 13-protofilament arrangement with an outer diameter of approximately 24 nm. (C, D) Section of the L. tarentolae microtubule showing four tubulin heterodimers, with the conserved M-loop (highlighted in yellow) forming lateral contacts with neighbouring tubulin heterodimers.

**Figure S3.**
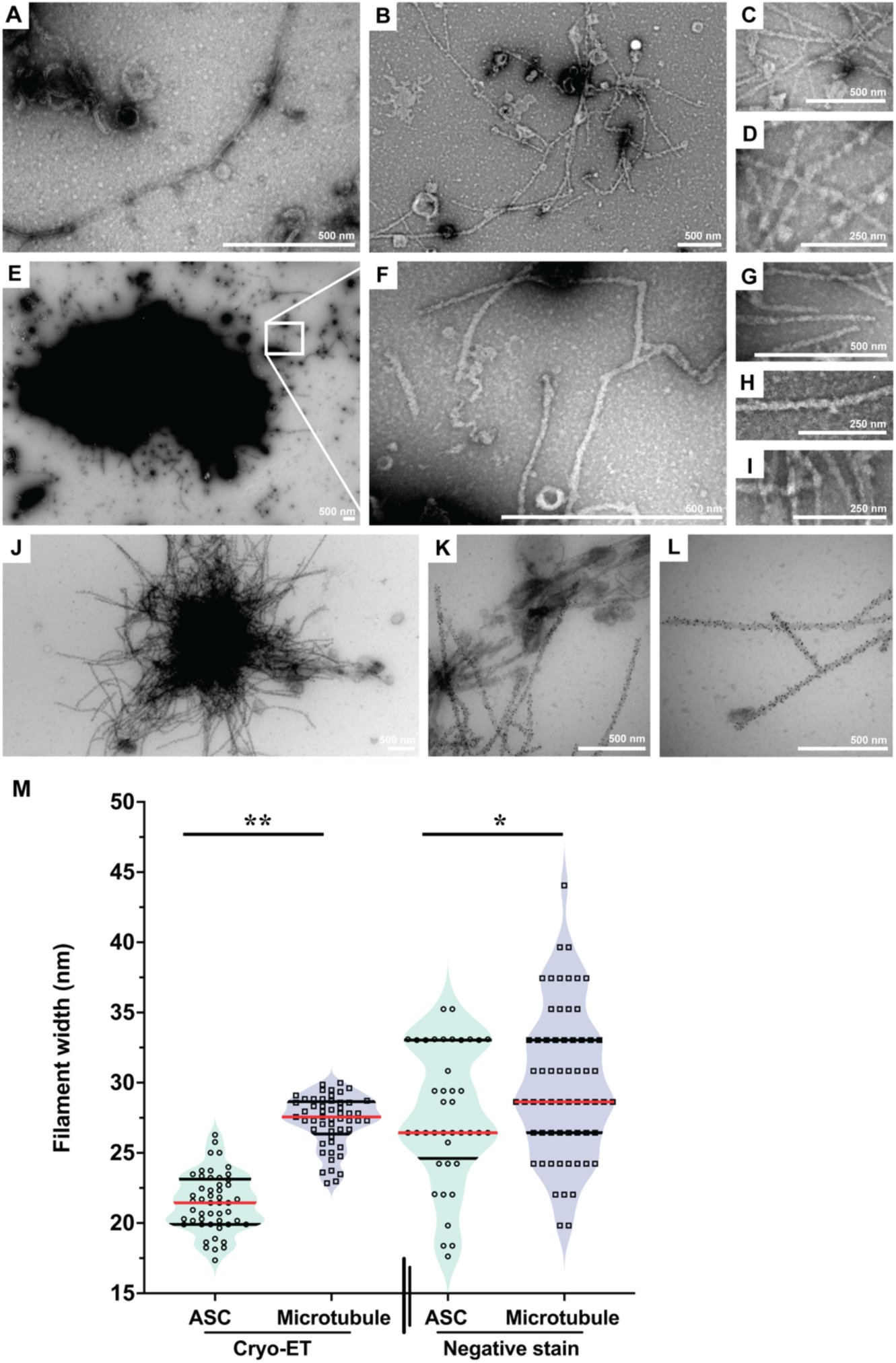
ASC^FL^-GFP cell-free expression. (A) Negative-stain EM of control lysate with no protein expression. (B-D) ASC^PYD^-untagged expression results in the formation of filaments throughout the lysate. (E-I) ASC^FL^-GFP expression generates filaments that extend from large electron-dense speck-like structures. (J-L) Immuno-EM confirming ASC^FL^-GFP specks are highly decorated with GFP. (M) Plot of filament width measurements for ASC^FL^-GFP and microtubules in cryo-ET and negative-stain Leishmania tarentolae lysate samples. Fifty measurements were performed per condition, with a maximum of three measurements per filament, spaced at least 100 nm apart along the filament length. Statistical significance is indicated as * p < 0.05 and ** p < 0.01.

**Figure S4.**
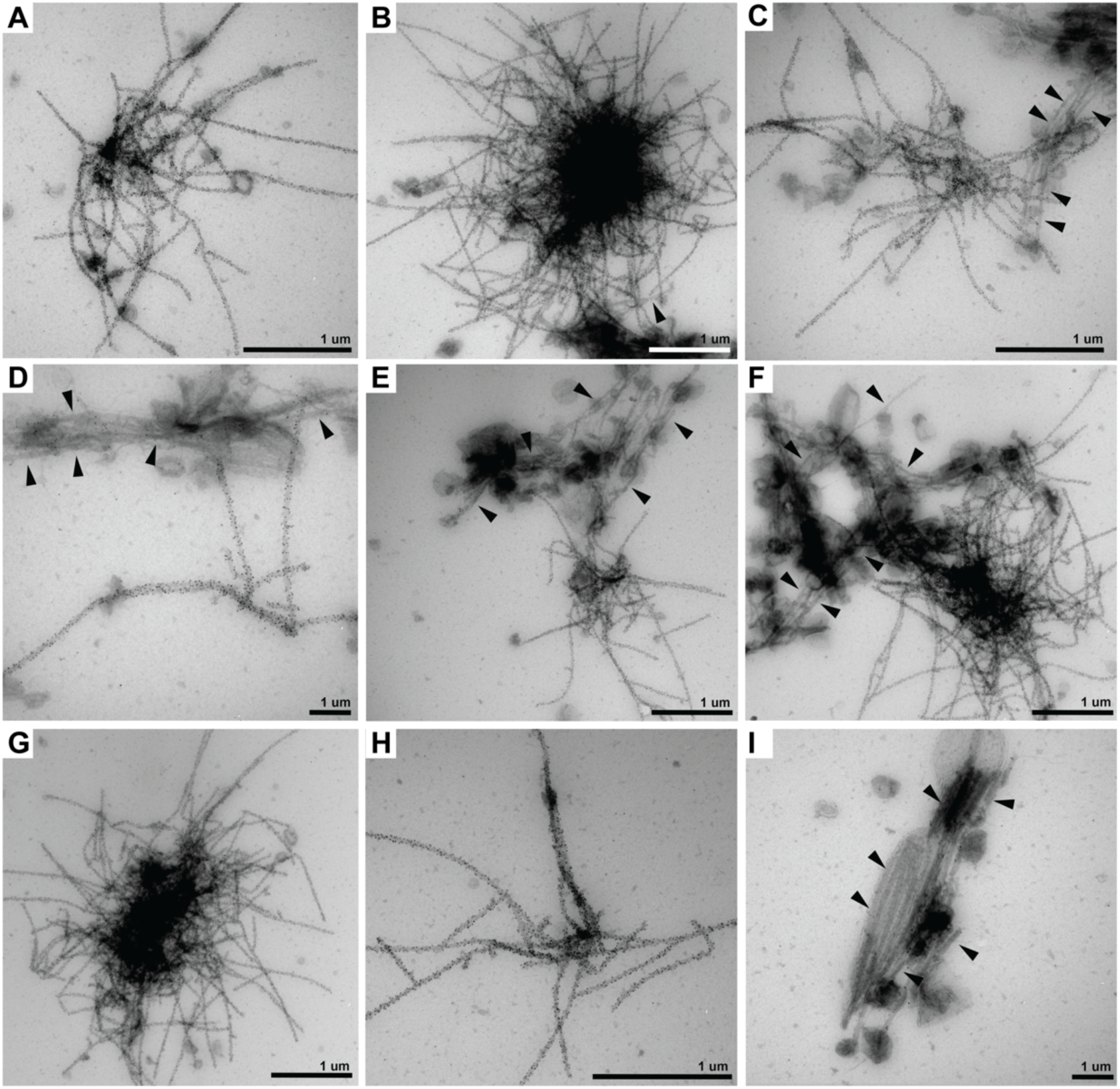
LTE expression of ASC^FL^-GFP generates structurally diverse complexes. (A-H) Immuno-gold labelled ASC^FL^-GFP, showing that the labelled filaments adopt a consensus structure with a core domain, from which filaments extend in all directions. The filaments extending from these central cores are regular in size and shape. 4.5nm anti-GFP immuno-gold was used. (I) No immunogold labelling of endogenous lysate microtubules shown by black arrow heads.

**Figure S5.**
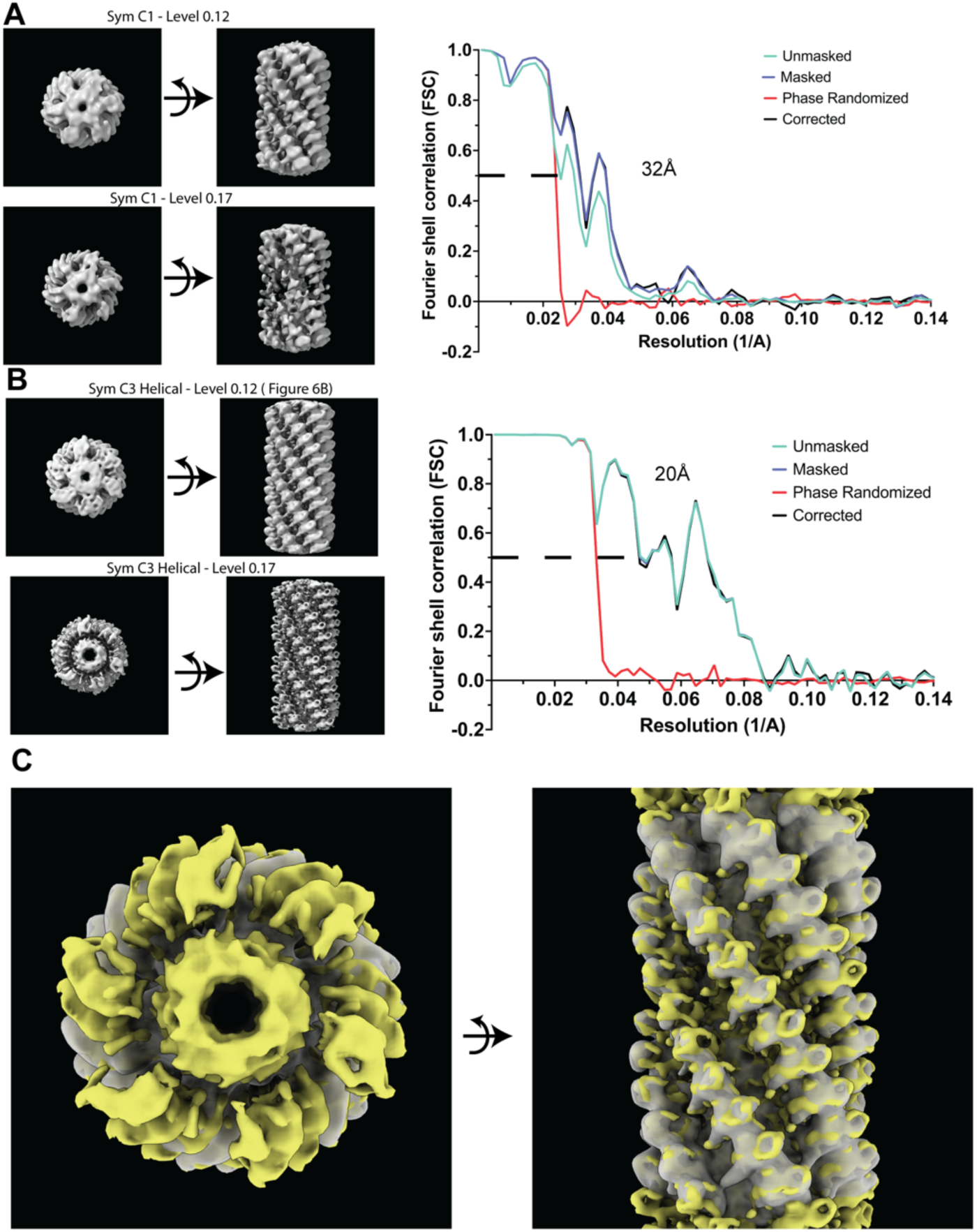
Sub-tomogram average reconstruction of ASC^FL^-GFP filaments. Sub-tomogram average reconstruction of ASC^FL^-GFP filaments (Class 1, 4724 particles) using different symmetry parameters: (A) symmetry C1; and (B) symmetry C3 with helical reconstruction, with differing level views in ChimeraX (Version 1.10) and the corresponding Fourier shell correlation (FSC) vs resolution (1/Å) plots, showing resolution estimates (FSC criterion 0.5). (C) Superimposed view of reconstructions (A and B), highlighting the helical reconstruction improving poorly resolved outer regions in C1 reconstruction. This dataset was acquired with a single defocus which has impacted the shape of the FSC curve.

**Figure S6.**
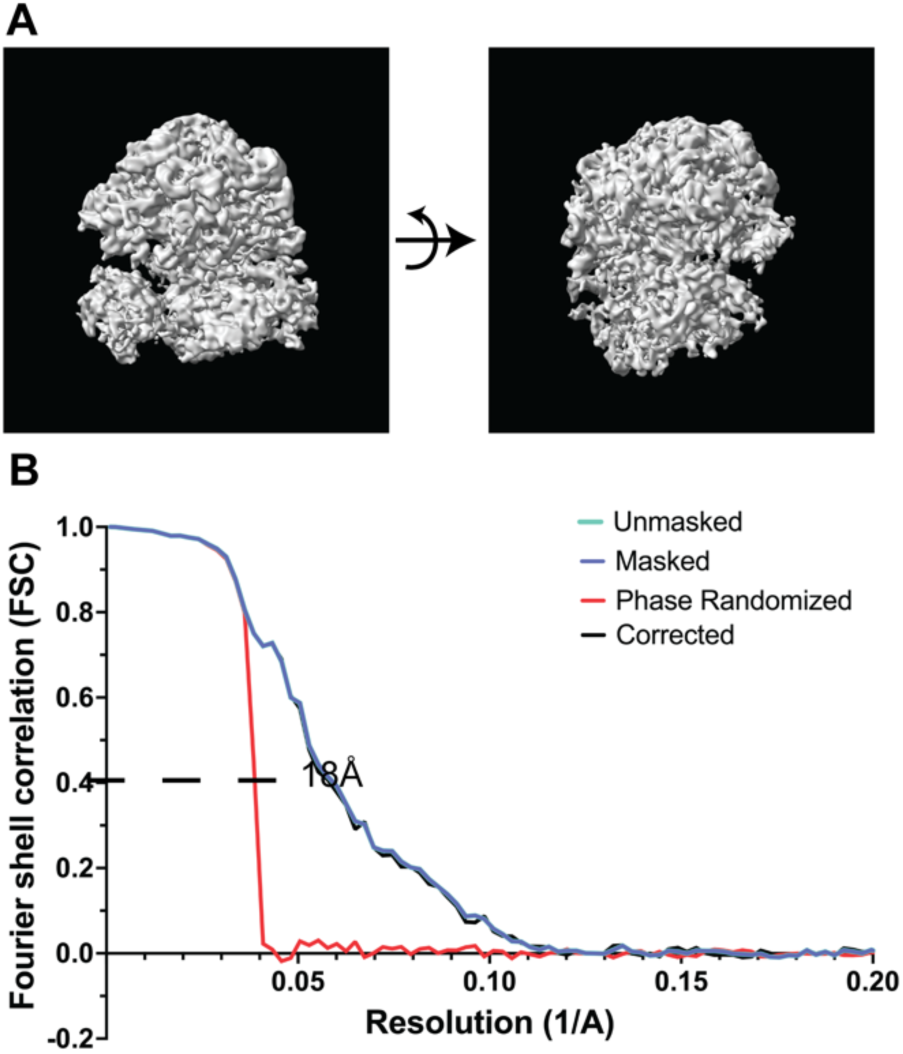
Sub-tomogram average structure of *L. tarentolae* ribosome, (A) showing the large and small subunit; and (B) the corresponding Fourier shell correlation (FSC) vs resolution (1/Å) plots, showing the resolution estimate of 29 Å (FSC criterion 0.5).

**Figure S7.**
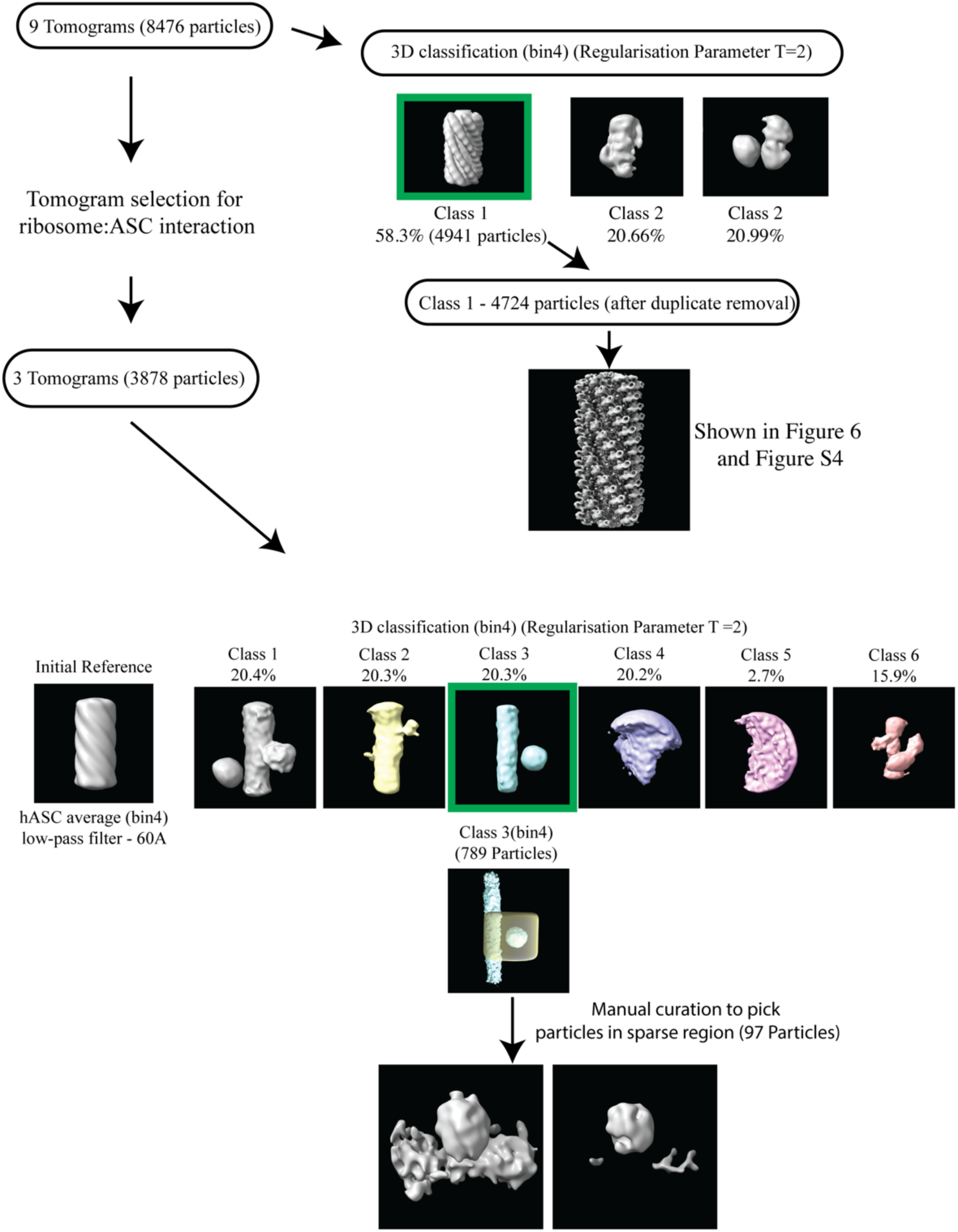
Structural and Heterogeneity Analysis of ASC^FL^-GFP Particles. A schematic illustrating the structural analysis of ASC^FL^-GFP filaments, starting from 9 tomograms (8,476 particles), followed by 3D classification to select Class 1 (4,724 particles) for sub-tomogram averaging, as shown in Figure 6A and 6B. A secondary pipeline for ASC:ribosome interaction analysis involves tomogram curation with selection of 3 tomograms and 3D classification to identify ribosome:ASC filament interactions, leading to the selection of Class 3 (marked in green) for further refinement, as shown in Figure 6D.

